# Structural Characterisation of Nanoparticle-Supported Lipid Bilayers by Grazing Incidence X-ray and Neutron Scattering

**DOI:** 10.1101/2022.07.07.499146

**Authors:** Nicolò Paracini, Philipp Gutfreund, Rebecca Welbourn, Juan Francisco Gonzalez, Kexin Zhu, Yansong Miao, Nageshwar Yepuri, Tamim A Darwish, Christopher Garvey, Sarah Waldie, Johan Larsson, Max Wolff, Marité Cárdenas

## Abstract

The structure of supported lipid bilayers formed on a monolayer of nanoparticles was determined using a combination of grazing incidence X-ray and neutron scattering techniques. Ordered nanoparticle arrays assembled on a silicon crystal using a Langmuir-Schaefer deposition were shown to be suitable and stable substrates for the formation of curved and fluid lipid bilayers that retained lateral mobility, as shown by fluorescence recovery after photobleaching. A comparison between the structure of the curved bilayer assembled around the nanoparticles with the planar lipid membrane formed on the flat underlying silicon oxide surface revealed a ∼5 Å thinner bilayer on the curved interface, resolving the effects of curvature on the lipid packing and overall bilayer structure. The combination of neutron scattering techniques, which grant access to sub-nanometre scale structural information at buried interfaces, and nanoparticle-supported lipid bilayers, offers a novel approach to investigate the effects of membrane curvature on lipid bilayers.

## Main text

Supported lipid bilayers (SLBs) are robust and widespread models of biological membranes used in applications ranging from biophysical studies of membrane function (1) to biosensing (2). Planar SLBs typically consist of phospholipid bilayers formed via vesicle fusion (3), solvent-assisted bilayer formation (4) or Langmuir-Blodgett and Langmuir-Schaefer monolayer transfer techniques (5,6) onto flat hydrophilic interfaces of materials like quartz, mica and silicon oxide. There is growing interest towards the development of non-planar SLBs that deviate from canonical flat interfaces and instead display a degree of curvature of the lipid bilayer. These model systems find applications in the study of membrane curvature-mediated phenomena such as lipid and protein segregation and binding of curvature-sensitive proteins (7–13). For this purpose, nano- and micropatterned surfaces represent attractive substrates for the formation of SLBs with well-defined surface topologies imparted by the underlying interfacial nanostructure.

Both top-down and bottom-up approaches have been adopted for the modification of flat interfaces and the formation of surface patterns suitable to form curved SLBs. Top-down methods are primarily based on nanolithography techniques which afford a high degree of control and fine tuning over the resulting surface structure (8,9). The high precision of top-down methods however often comes at the cost of time and resources, which can become limiting factors when dealing with large surfaces and number of substrates to functionalise. Bottom-up approaches, on the other hand, typically rely on self-assembly processes driven by chemical and physical forces and can be exploited to fabricate patterned samples using nanoparticles (NP) to serve as substrates for SLB formation (7,10,14). Amongst bottom-up methods employed to form large arrays of NPs, Langmuir-Blodgett and Langmuir-Schaefer depositions offer an additional level of control on the self-assembly process by enabling the adjustment of the packing density of the Langmuir monolayer at the air/water interface prior to its transfer onto a solid substrate (15–17). Furthermore, Langmuir transfer techniques have the advantage of yielding large uniform monolayers which make well-suited samples for characterization by flux-limited grazing incidence scattering methods such as neutron reflectometry (NR) and grazing incidence neutron small angle scattering (GISANS) (18,19). Due to their unique ability to probe non-invasively buried interfaces and their differential sensitivity towards hydrogen and deuterium, neutrons are amongst the most powerful surface sensitive techniques for the structural characterization of complex biological thin films at solid/liquid interfaces, of which SLBs represent a primary example. Grazing incidence neutron scattering, particularly NR, has found wide application in the structural characterisation of planar SLBs, however the potential of techniques like GISANS, as well as NR, remains largely untapped when it comes to structural studies of model membranes with more complex topologies. In this context NR and GISANS, combined with selective lipid deuteration, can provide a novel approach to study the effect of curvature on the in-plane and out-of-plane structural features of SLBs.

In this article, we exploit a modified Langmuir-Schaefer deposition method to form large arrays of spherical silica NP on silicon oxide surfaces that are used as substrates to form nanoparticle-supported lipid bilayers (NP-SLB). First, we combine X-ray and neutron scattering to probe the structure of the NP arrays in both dry and aqueous environments (i.e. at the solid/air and solid/liquid interface). In-plane and out-of-plane structural properties of the NP-SLB system are resolved by NR and GISANS. Finally, using fluorescence recovery after photobleaching we show that lipids in the NP-SLB retain lateral mobility.

The formation of densely packed arrays of non-porous silica NPs onto a polished silicon substrate was achieved via a modified Langmuir-Schaefer transfer of NPs from the air/water interface onto a submerged substrate (20,21) as depicted in **Figure 1A**. To maximise the particle density in the monolayer, pressure-area isotherms were recorded to establish the maximum surface pressure applicable to the NP monolayer at the air-water interface before excessive compression resulted in a collapse, indicated by an abrupt change in surface pressure (**Figure 1B**). A value slightly below the collapse pressure was selected for NP depositions. Atomic force microscopy (AFM) confirmed the formation of a NP layer with high density of particles packed in a hexagonal lattice and interspersed with minor defects (**Figure 1C**).

**Figure 1.**
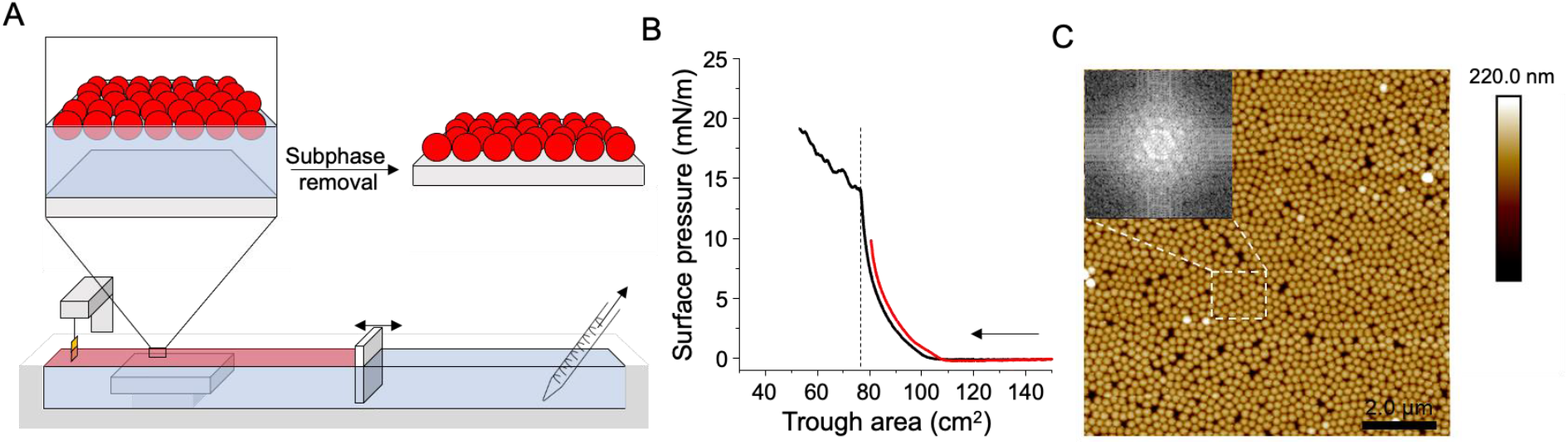
Langmuir-Schaefer transfer of a NP monolayer onto a solid substrate. (**A**) Schematic representation of the modified Langmuir-Schaefer transfer of the silica NP monolayer onto a submerged silicon crystal. Once the NP monolayer (red area) was compressed to the target surface pressure, measured by the Wilhelmy plate, the aqueous subphase was slowly removed from behind the barrier using a serological pipette tip connected to a pump. (**B**) Pressure-area isotherm of NP monolayer compressed above (black) and below (red) the collapse point at 15 mN/m and indicated by an abrupt change in the slope (dashed line), the arrow indicates the direction of compression. (**C**) NP monolayer (nominal diameter 2000 Å) after transfer onto a silicon wafer imaged by AFM. Black areas correspond to gaps between the particles whilst bright spots are particles absorbed on top of the NP monolayer. The scale bar is 2 µm. The inset shows a Fourier transform of the highlighted region in the AFM image and the corresponding hexagonal lattice.

Imaging techniques, such as AFM, only provide local information over small areas. Scattering methods on the other hand, probe the average structure over large areas in both dry and aqueous environments. Following the assembly method illustrated in **Figure 1**, monolayers of commercial NP with nominal diameters of 50, 100 and 200 nm were characterized by NR and GISAXS at the solid/liquid and air/solid interfaces. (**Figure 2**). The samples, assembled into custom built solid/liquid cells, were first characterised by NR in an aqueous environment. NR measurements enable the determination of the structure of buried thin films along the axis perpendicular to the substrate, yielding a profile describing the scattering length density (SLD) distribution, which determines the neutron refractive index along the normal to the surface. Reflectivity curves measured in the presence of H_2_O and D_2_O were fitted simultaneously to a model of the interface describing a monolayer of spheres (**Figure S1**).

**Figure 2.**
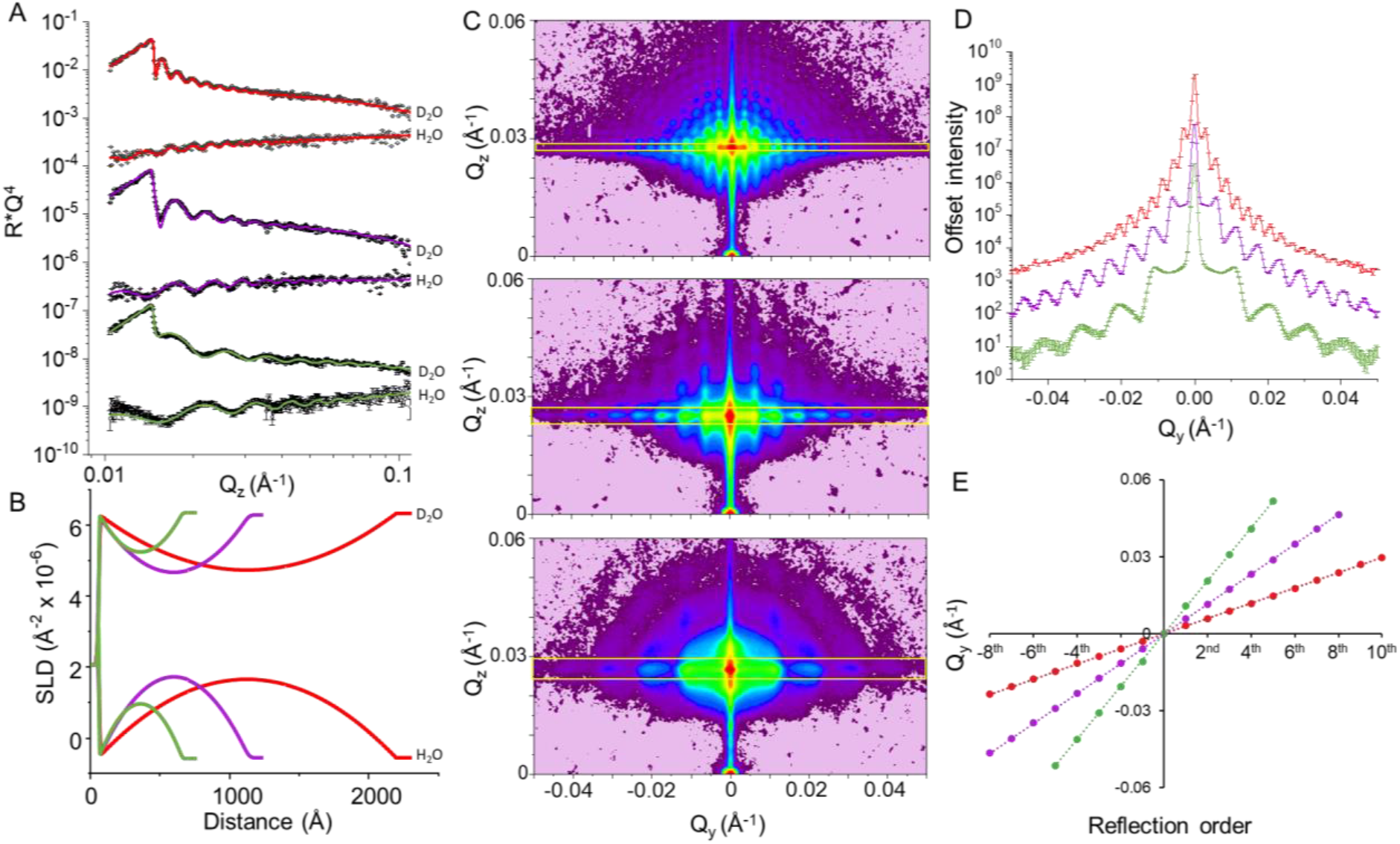
Structural characterization of NP monolayers by NR and GISAXS. (**A**) Neutron reflectometry data (points) and best fit (lines) of three monolayers containing NP of different sizes, measured in D_2_O and H_2_O. Data and corresponding fits are offset vertically for clarity (**B**) corresponding SLD profile derived from the constrained fit of NR data from the two solution isotropic contrasts for each NP set. (**C**) GISAXS detector images of three monolayers measured at the air/solid interface (**D**) Plot of the integrated intensities within the yellow boxes shown in **C**. Absolute intensity is offset for clarity (**E**) Linear regressions of the maxima positions along *Q*_*y*_ of the peaks shown in **D**.

The SLD profiles obtained from the constrained fits to the reflectivity curves collected in H_2_O and D_2_O (see methods and **Figure S4** for details on the modelling), yielded a NP monolayer thickness of 2149 ± 12 Å, 1086 ± 8 Å and 604 ± 5 Å for the three samples, in good agreement with the size of the particles, confirming the formation of NP monolayers (**Figure 2A**). From the contrast provided by the hydrogenous and deuterated water the in-plane packing density of the particles in the monolayers was extracted, ranging from 66 ± 1 % for the largest particles, 62 ± 5 % for the intermediate size to 40 ± 4 % for the smallest NPs (**Table S1**). A value of 100% corresponds to a defect-free layer of spheres tightly packed in a hexagonal lattice. The significantly worse packing of the smallest particles is indicative of larger areas occupied by defects in the NP monolayer, which might result from a size-dependent tendency to form inhomogeneous aggregates during the Langmuir-Schaefer procedure.

To complement the information obtained by NR, the samples were characterised by GISAXS which is sensitive to the in-plane arrangement and correlations between the NPs. The three particle sizes gave rise to a strong GISAXS signal displaying several orders of well-defined peaks (**Figure 2C**). The images were integrated across the *Q*_*y*_ axis and over a *Q*_*z*_ range corresponding to the angle of specular reflection and adjusted to include a single row of peaks in each integration box in order to extrapolate the in-plane correlation distances related to the patterns observed (**Figure 2D**, integrated regions shown by the yellow boxes in **Figure 2C**). The linear plot of the *Q*_*y*_ position of the maxima against the order of the peaks indicates that the maxima are equidistant in *Q*_*y*_, and the slope of the straight lines fitted through the data points corresponds to the average Δ*Q*_*y*_ (**Figure 2E**). According to the inverse relationship between the *Q*_*y*_ spacing of the maxima and the corresponding real space distances, given by *d=2π*/ Δ*Q*_*y*_, the signals from the different samples yielded correlation distance values of 2124 Å, 1081 Å and 609 Å. The parameters obtained from the model-free analysis of the GISAXS patterns were found to be in close agreement with the monolayer thickness values obtained from the NR fits. The information on particles density and size obtained from NR were used as input parameters to generate simulated GISAXS signals using the distorted wave Born approximation (DWBA) implemented in the BornAgain software (22). The simulated scattering was in good agreement with the data, providing qualitative information on the average interparticle distances and on the degree of order in the monolayers (**Figure S2**). To obtain simulations that reproduced the intensity and *Q*_*y*_ spacing of the data, an increasing level of disorder of the in-plane NP monolayer structure had to be assumed as particle sizes decreased, which is consistent with the lower packing densities extracted from the NR model.

The monolayers containing the largest NP displayed the highest level of order as well as the highest particle density and were therefore chosen as the substrate for the study of lipid bilayer deposition. Lipid deposition via vesicle fusion was monitored by quartz crystal microbalance with dissipation (QCMD) which was used to compare the process of bilayer formation on conventional “flat” silicon oxide sensors and on sensors coated with the NP monolayer (**Figure 3A**). The frequency shift (Δ*f*) observed after injecting POPC vesicle and rinsing with water was −24.2 ± 0.7 Hz and −25.5 ± 0.8 Hz on the flat and NP-coated sensors respectively, in line with the values reported previously for the formation of lipid bilayers on flat silicon oxide sensors as well as on QCMD sensors coated with NP (7). The shift in dissipation (Δ*D*) amounted to +0.4 ± 0.3 ppm on the flat and −5.8 ± 1.2 ppm on the NP-coated surfaces (**Figure S3**). The slightly higher Δ*f* value recorded on the NP sample shows a marginally higher amount of lipids adsorbed whilst the negative Δ*D* indicates that lipid addition affects the properties of the NP layer, suggesting that intercalation of lipids in between the NP increases the overall stiffness of the NP-SLB layer (23). The surfaces were then rinsed with ethanol to remove the POPC which in both cases caused Δ*f* and Δ*D* to return to the baseline measured at the beginning of the experiment, indicating complete removal of the lipids by the ethanol wash, with the NP-array remaining unperturbed. A second cycle of lipid deposition and removal was carried out yielding Δ*f* and Δ*D* values in line with the shifts observed in the first cycle, showing full reusability of the NP-array as a substrate for the formation of NP-SLBs (**Figure 3A**).

**Figure 3.**
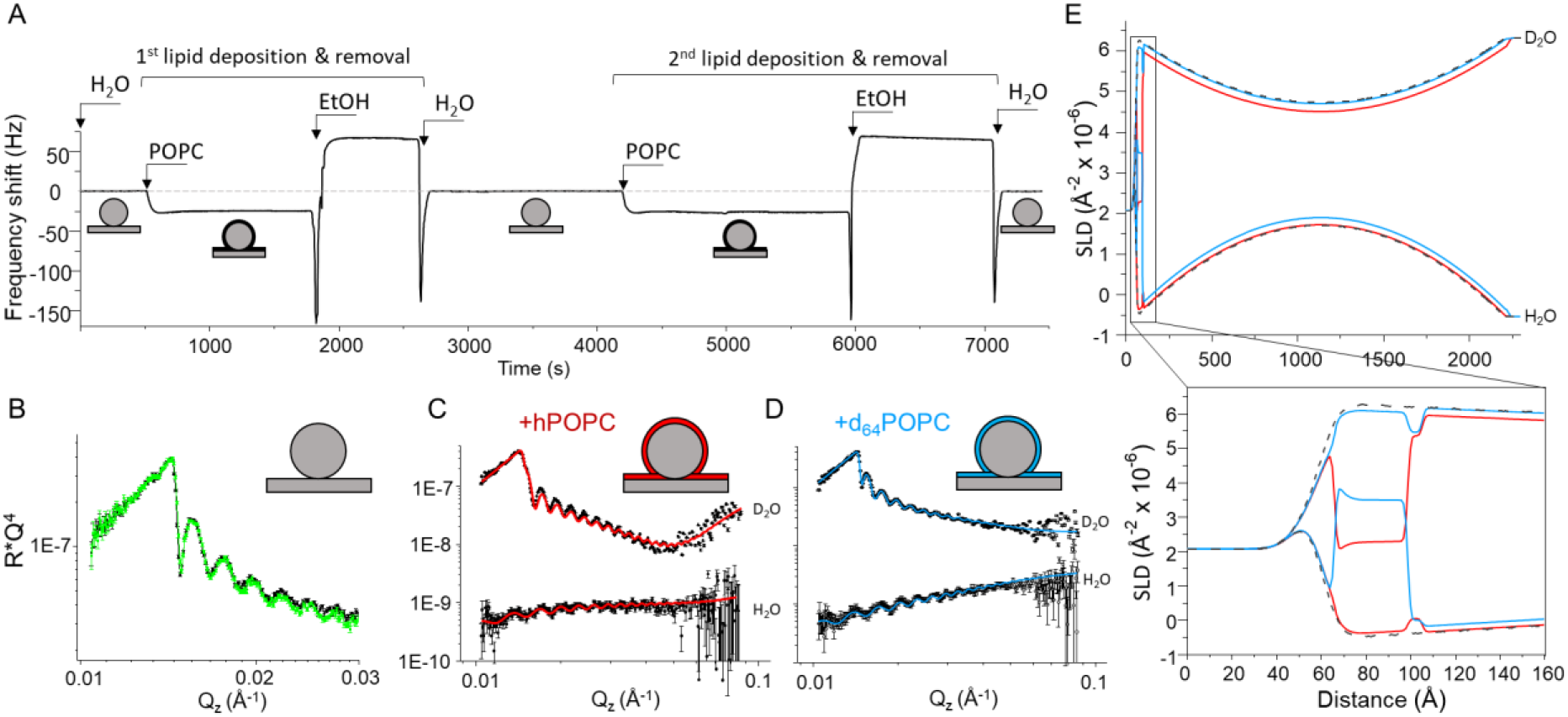
Formation of NP-SLB and vertical structure measured by NR. (**A**) QCMD trace monitoring the deposition of POPC onto a silica sensor coated with a monolayer of NP. Labels indicate the injection of different solutions in the QCMD flow module. The light grey flat dotted line is a guide to the eye set at zero Hz frequency shift. (**B**) NR profiles of NP monolayer in D_2_O before (black) and after (green) one cycle of lipid deposition and removal, showing recovery of the reflectometry signal upon EtOH rinsing (log-log scale) (**C**) NR data (points) and best fit (red lines) of the NP-SLB formed with hPOPC in H_2_O and D_2_O. (**D**) NR data (points) and best fit (blue lines) of the NP-SLB formed with d_64_POPC in H_2_O and D_2_O (**E**) SLD profiles describing the NP monolayer in the absence of lipids (black dashed lines) in the presence of hPOPC (red lines) and in the presence of d_64_POPC (blue lines). The expanded region shows a magnification of the flat SiO_2_ interface highlighting the formation of a planar SLB on the flat silicon substrate.

The NP-SLB was characterised by NR to access information on the structure of the lipid bilayer formed on the NP array. To fully exploit the ability of neutrons to differentiate between hydrogenous and deuterated molecules, the process investigated by QCMD was replicated on the neutron beam line using first hydrogenous hPOPC, followed by regeneration of the surface with ethanol and deposition of tail deuterated d_64_POPC. After each lipid assembly, the sample was characterised both in H_2_O and in D_2_O. The reflectivity data sets collected in H_2_O and D_2_O on the bare NP array, the hydrogenous NP-SLB and the deuterated NP-SLB were then fitted to a common model of the interface where the structural parameters describing the NP monolayer were shared across the three conditions whilst the SLD of the lipids and their volume fractions were allowed to vary (see **Figure 2D, E** for the fits and SLD of the NP monolayer before lipids addition). The model that produced the most accurate fit to the reflectivity data from the NP-SLB samples contained a lipid bilayer coating the entire nanoparticle surface as well as part of the flat surface of the silicon substrate supporting the particles (See **Figure S4** for a detailed description of the model used). Addition of hPOPC vesicles to the NP array caused a prominent shift of the reflectivity profile measured in D_2_O whilst leaving the signal recorded in H_2_O mostly unaffected, as expected from the deposition of hydrogenous material at the interface. Conversely, addition of d_64_POPC resulted in a large shift in the reflectivity measured in H_2_O but no significant changes in D_2_O (**Figure S5**). Similarly to what was observed with QCMD, rinsing the NP-SLB with ethanol between bilayer depositions reverted the reflectivity back to the initial profile measured before lipid addition, confirming a nearly complete removal of the lipids whilst preserving the intact NP monolayer structure (**Figure 3B**).

The parameters obtained from the fits to the NR data described the formation of a lipid bilayer adhering to the NP, which coated the entire surface of the spheres with a high coverage as indicated by the low volume fraction of water detected in the tail region, below 4% in both the hPOPC and d_64_POPC hydrophobic cores (**Table S2, Figure S6**). Additionally, a planar lipid bilayer formed on the silicon crystal surface underlying the NP monolayer, with an overall coverage of ∼60%, indicating the formation of a less complete lipid layer in comparison with the bilayer coating the NP. The lower coverage of the underlying planar SLB is likely to result from the partial shadowing by the large NP which prevented the formation of a homogeneous continuous lipid layer (**Figure S4A**). Along with the differences in coverage, the structural data revealed that the NP-SLB displayed a ∼27 Å thick hydrophobic core, ∼5 Å thinner than the planar SLB (∼32 Å), as well as a ∼20% higher water content associated with the lipid head groups. Whilst the parameters obtained for the planar POPC SLB are in excellent agreement with previously reported values (24,25), the thinner structure of the NP-SLB pointed to a sub-optimal packing of the lipids around the spheres resulting from the curved substrate. Mean molecular area values, calculated assuming POPC tails molecular volume of 925 Å^3^ (24) yield values of ∼58 Å^2^ and ∼69 Å^2^ for the planar and curved SLB respectively, corroborating the looser packing of POPC in the latter.

The NR curves measured in D_2_O showed a strong off-specular signal captured in the 2D detector images as horizontal stripes in the region of total reflection (**Figure 5A**) which caused the intensity dips visible in the specular reflectivity signal below the critical edge (**Figure S7**). This is similar to what is observed in the case of resonators, characterised by a potential well, formed of a region of low SLD in between two regions of high SLD (26). In the case of the spherical NP system under investigation here, the increasing volume fraction of silicon oxide (SLD of SiO_2_ = 3.47×10^−6^ Å^-2^) towards the centre of the NP monolayer generates a region of low SLD in comparison to the D_2_O rich regions above and below the monolayers centre (SLD of D_2_O = 6.35×10^−6^ Å^-2^), which may explain the appearance of the resonances observed.

The addition of hPOPC caused both an increase in the intensity of the off-specular scattering and a shift in the *Q*_*z*_ position of the resonances (**Figure 4A**). Simulations of the off-specular data (27) reproduced qualitatively the signal change observed for the NP monolayer before and after addition of lipids (**Figure 4B**) confirming the validity of the specular reflectivity model used to fit the data. Moreover, the off-specular scattering simulations supported the GISAXS results indicating better ordering in the larger NP arrays. In order to reproduce the measured off-specular intensities of the 100 nm and 50 nm particles (**Figure S8**), micrometre sized D_2_O clusters had to be included in between the NPs in the calculations, whilst these large D_2_O clusters were not required to simulate the signal from the 200 nm particles.

**Figure 4.**
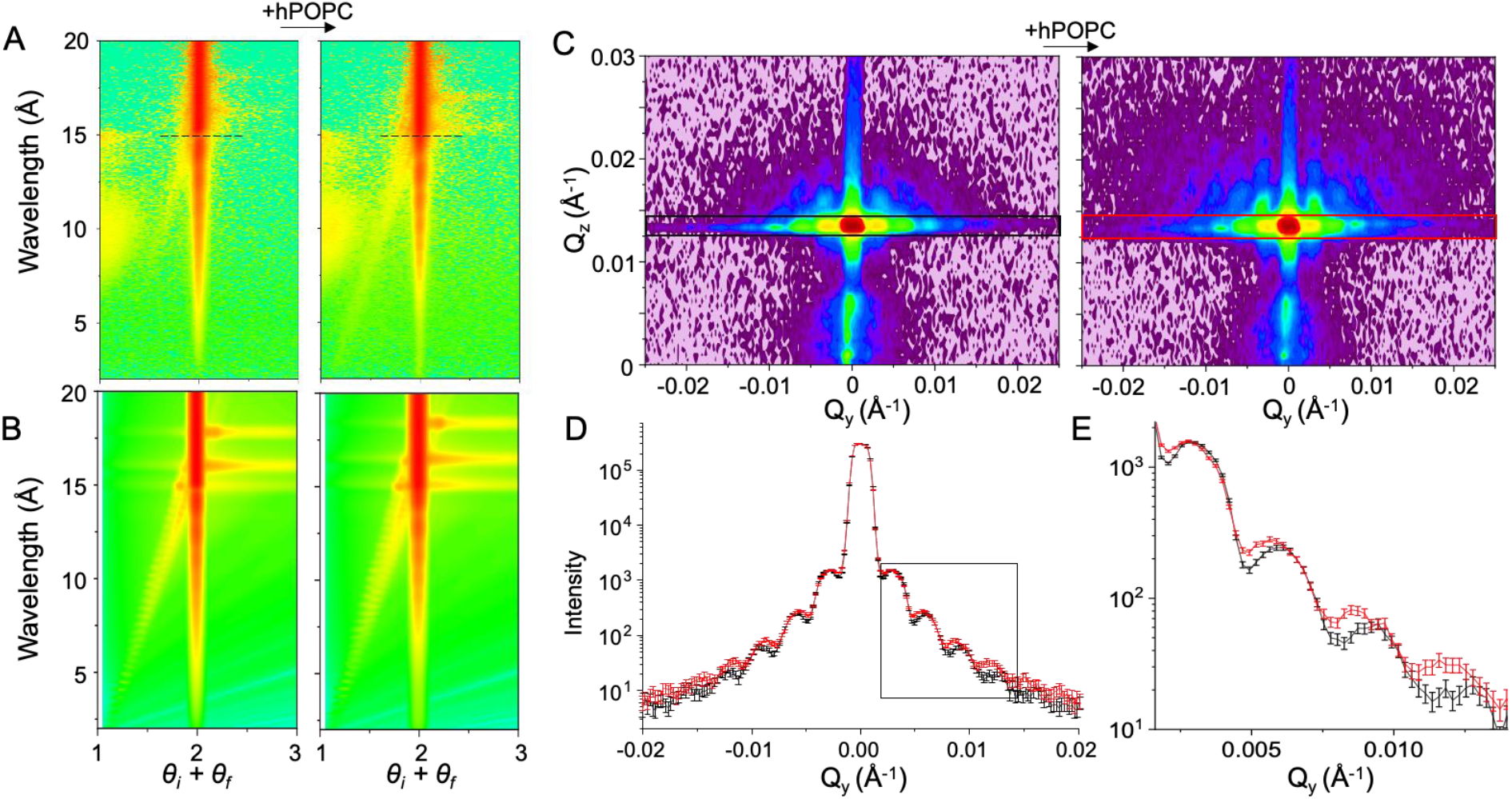
NP-SLB formation monitored by GISANS and off-specular NR. (**A**) Off-specular NR data in D_2_O before (left) and after (right) the addition of hPOPC to the NP-array (dashed lines indicate the position of the critical edge) (**B**) Simulations of off-specular NR in the absence (left) and presence (right) of lipids coating the NP *θ*_*i*_ and θ_*f*_ are the incident and reflected angles respectively (**C**) GISANS detector images of NP monolayer in D_2_O before (left) and after (right) the addition of hPOPC (**D**) Integration across *Q*_*y*_ of the GISANS images corresponding to the coloured boxes in **C** before (black) and after (red) addition of lipids (**E**) Magnified region of the peaks marked in **D**

Following up on the GISAXS data collected on the dry NP arrays, the process of NP-SLB formation was investigated by GISANS, which enables measurements of grazing incidence small angle scattering on buried (e.g. wet) biological thin films. The analysis of the integrated peak positions yielded a correlation distance of 2083 Å prior to the addition of lipids, in good agreement with the values obtained from the GISAXS and NR results (**Figure 2**). A comparison of the GISANS signals before and after addition of hPOPC to the NP array in D_2_O revealed an increase in the intensity of the overall scattering in the 2D detector image, in line with the increased contrast in the sample caused by the addition of hydrogenous lipids in D_2_O (**Figure 4C, D**). Notably, along with the change in intensity, the adsorbed lipids caused a shift of the maxima positions in *Q*_*y*_, with a reduction of the average inter-peak separation, resulting in an increase in the correlation distance from 2083 Å to 2167 Å, corresponding to an overall increase of 84 Å in the particle diameter, as calculated form the linear regression of the peaks positions (**Figure S9**).

According to the structural information obtained from the NR data the total bilayer thickness formed around the NP amounts to ∼39 Å, thus the expected increase in the apparent NP diameter upon bilayer formation would be ∼78 Å, in good agreement with the 84 Å obtained from the model-free GISANS analysis which corroborated the NR results.

Finally, diffusivity of the POPC molecules within the NP-SLB was investigated using FRAP. When compared to SLB formed onto flat glass surfaces, the measured lipid diffusion after bleaching was slower on the bilayers assembled on the NP coated substrate (**Figure 5A**). The final recovery was nonetheless comparable between the two substrates after 120s from the bleaching, indicating that lipid mobility in the NP-SLB, although more restricted, was still largely retained both in the planar bilayer and around the NP (**Figure 5B**). Further experiments are required to understand to what extent diffusion takes place directly between neighbouring NP compared to diffusion mediated by the planar underlying SLB. Imaging of the NP-SLB by super resolution (SR) and total internal reflection fluorescence (TIRF) microscopy provided a clear picture of the fluorescent bilayer and the NP array. SR and TIRF images were acquired on NP-SLB formed on particles with a larger nominal diameter (400 nm) that provide the optimal resolutions for the SLBs at different Z-positions, including the NP-SLB and the planar SLB on the underlying glass surface (**Figure 5C**).

**Figure 5.**
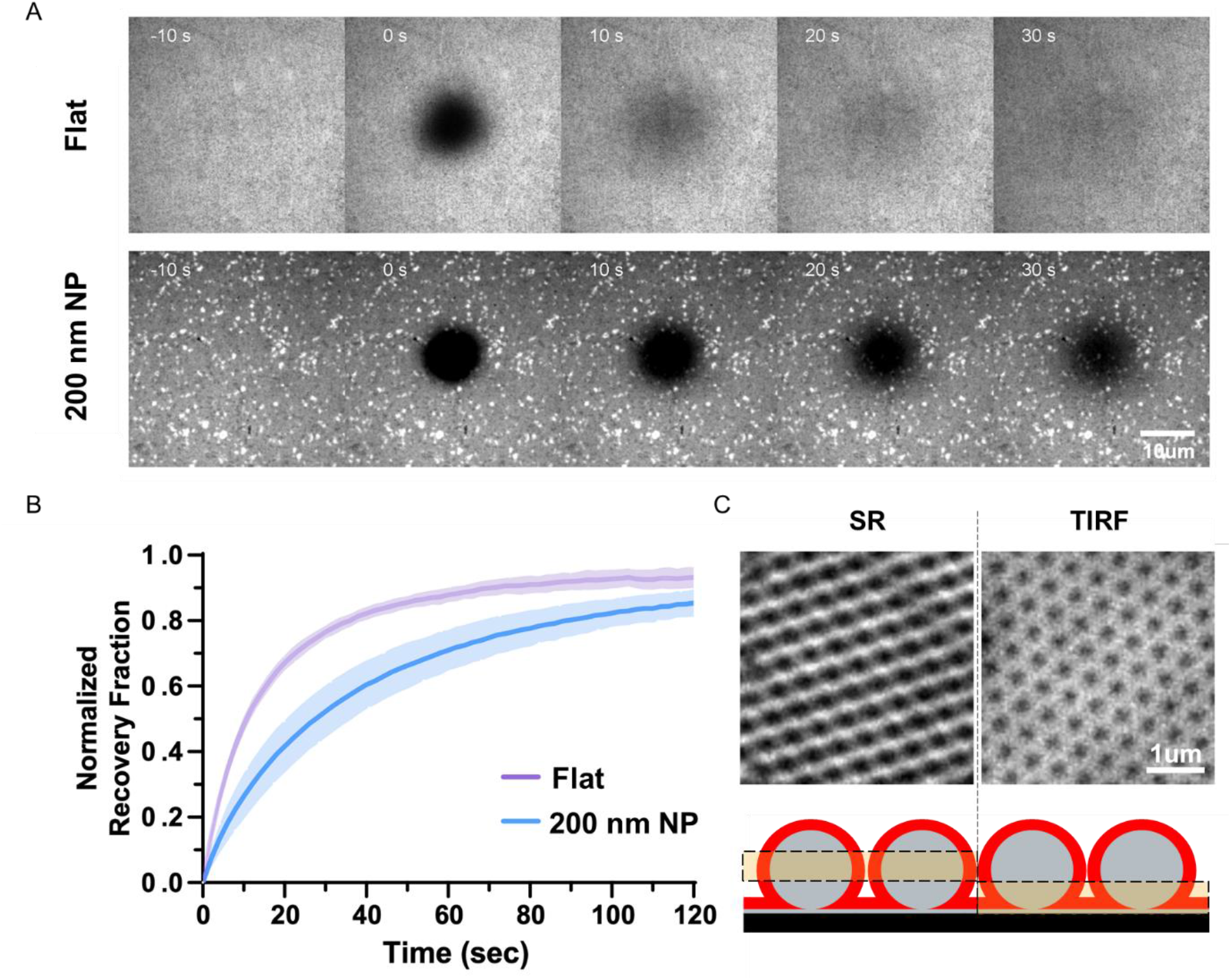
Lipid mobility measured by FRAP. (**A**) FRAP images of POPC bilayers deposited on flat (top) and substrates coated with 200 nm NP, the scale bar is 10 µm. (**B**) Recovery of the fluorescence intensity after photobleaching showing the average of different measurements from five different region of interest with the standard deviation represented by the shaded area. (**C**) Super resolution (SR) microscopy and total internal reflection fluorescence (TIRF) images of NP array coated with a POPC bilayer; the highlighted boxes in the cartoon show the regions where each fluorescent microscopy technique is most sensitive. In SR and TIRF imaging experiments NP with a larger nominal diameter (400 nm) were used in order to obtain a clearer image of the sample.

The properties of NP-SLBs have been investigated both in bulk (28–32) and at interfaces (7,10,14,33) with a wide range of biophysical techniques. Here, for the first time, we provide a close up on the molecular structure of the lipid bilayer assembled on nanoparticles using a combination of surface sensitive techniques. Together the data demonstrate the possibility of using large NP arrays assembled via an accessible bottom-up method that does not involve complex nanofabrication, as substrates for the formation of high-coverage curved lipid bilayers. The combination of scattering and imaging methods provides access to accurate structural information on the vertical and in-plane structure of the NP-SLB. The NP-SLB platform described here provides a new tool that can be employed in the study of curvature dependent phenomena using both grazing incidence scattering and imaging techniques.

## Acknowledgements

MC and NP thank the Swedish Research Council for financial support. NP acknowledges support from Nordforsk - Nordic Neutron Science Program (Grant 106881). MC thanks Wennergren foundation for financial support. The authors thank the ILL for beamtime (D22: doi:10.5291/ILL-DATA.8-02-912, FIGARO: doi:10.5291/ILL-DATA.8-02-889). The authors thank Prof. Jaume Torres for access to a Langmuir Trough at Nanyang Technological University. This study was supported by Singapore MOE Tier 3 (MOE2019-T3-1-012) to Y.M. The National Deuteration Facility in Australia is partly funded by The National Collaborative Research Infrastructure Strategy (NCRIS), an Australian Government initiative.

## Abbreviations

NP: nanoparticles
SLB: supported lipid bilayer
SLD: scattering length density
NR: neutron reflectometry
GISAXS: grazing incidence small angle X-ray scattering
GISANS: grazing incidence small angle neutron scattering
FRAP: fluorescence recovery after photobleaching

## Notes

### Competing Interest Statement

The authors have declared no competing interest.

## References

1. Clifton LA, Campbell RA, Sebastiani F, Campos-Terán J, Gonzalez-Martinez JF, Björklund S, et al. Design and use of model membranes to study biomolecular interactions using complementary surface-sensitive techniques. Adv Colloid Interface Sci. 2020;277.

2. Santoro F, Lubrano C, Matrone GM, Iaconis G. New frontiers for selective biosensing with biomembrane-based organic transistors. ACS Nano. 2020 Oct 27;14(10):12271–80.

3. Hardy GJ, Nayak R, Zauscher S. Model cell membranes: Techniques to form complex biomimetic supported lipid bilayers via vesicle fusion. Curr Opin Colloid Interface Sci [Internet]. 2013 Oct;18(5):448–58. Available from: https://linkinghub.elsevier.com/retrieve/pii/S1359029413000915

4. Tabaei SR, Choi J-H, Haw Zan G, Zhdanov VP, Cho N-J. Solvent-Assisted Lipid Bilayer Formation on Silicon Dioxide and Gold. Langmuir [Internet]. 2014 Sep 2;30(34):10363–73. Available from: http://www.ncbi.nlm.nih.gov/pubmed/25111254

5. Kurniawan J, Ventrici De Souza JF, Dang AT, Liu GY, Kuhl TL. Preparation and Characterization of Solid-Supported Lipid Bilayers Formed by Langmuir-Blodgett Deposition: A Tutorial. Langmuir. 2018;34(51):15622–39.

6. Paracini N, Clifton LA, Skoda MWA, Lakey JH. Liquid crystalline bacterial outer membranes are critical for antibiotic susceptibility. Proc Natl Acad Sci [Internet]. 2018;201803975. Available from: http://www.pnas.org/lookup/doi/10.1073/pnas.1803975115

7. Sundh M, Svedhem S, Sutherland DS. Formation of Supported Lipid Bilayers at Surfaces with Controlled Curvatures: Influence of Lipid Charge. J Phys Chem B [Internet]. 2011 Jun 23;115(24):7838–48. Available from: https://pubs.acs.org/doi/10.1021/jp2025363

8. Hsieh W-T, Hsu C-J, Capraro BR, Wu T, Chen C-M, Yang S, et al. Curvature Sorting of Peripheral Proteins on Solid-Supported Wavy Membranes. Langmuir [Internet]. 2012 Sep 4;28(35):12838–43. Available from: https://pubs.acs.org/doi/10.1021/la302205b

9. Parthasarathy R, Yu C, Groves JT. Curvature-Modulated Phase Separation in Lipid Bilayer Membranes. Langmuir [Internet]. 2006 May 1;22(11):5095–9. Available from: https://pubs.acs.org/doi/10.1021/la060390o

10. Black JC, Cheney PP, Campbell T, Knowles MK. Membrane curvature based lipid sorting using a nanoparticle patterned substrate. Soft Matter [Internet]. 2014;10(12):2016–23. Available from: https://www.rsc.org/softmatter

11. Vasilca V, Sadeghpour A, Rawson S, Hawke LE, Baldwin SA, Wilkinson T, et al. Spherical-supported membranes as platforms for screening against membrane protein targets. Anal Biochem [Internet]. 2018 May;549:58–65. Available from: https://doi.org/10.1016/j.ab.2018.03.006

12. Sanii B, Smith AM, Butti R, Brozell AM, Parikh AN. Bending Membranes on Demand: Fluid Phospholipid Bilayers on Topographically Deformable Substrates. Nano Lett [Internet]. 2008 Mar 1;8(3):866–71. Available from: https://pubs.acs.org/doi/10.1021/nl073085b

13. Dabkowska AP, Niman CS, Piret G, Persson H, Wacklin HP, Linke H, et al. Fluid and Highly Curved Model Membranes on Vertical Nanowire Arrays. Nano Lett [Internet]. 2014 Aug 13;14(8):4286–92. Available from: https://pubs.acs.org/doi/10.1021/nl500926y

14. Roiter Y, Ornatska M, Rammohan AR, Balakrishnan J, Heine DR, Minko S. Interaction of nanoparticles with lipid membrane. Nano Lett. 2008;8(3):941–4.

15. Vegso K, Siffalovic P, Jergel M, Weis M, Benkovicova M, Majkova E, et al. Silver Nanoparticle Monolayer-to-Bilayer Transition at the Air/Water Interface as Studied by the GISAXS Technique: Application of a New Paracrystal Model. Langmuir [Internet]. 2012 Jun 26;28(25):9395–404. Available from: https://pubs.acs.org/doi/10.1021/la301577a

16. Wu L, Wang X, Wang G, Chen G. In situ X-ray scattering observation of two-dimensional interfacial colloidal crystallization. Nat Commun [Internet]. 2018 Dec 6;9(1):1335. Available from: http://dx.doi.org/10.1038/s41467-018-03767-y

17. Guo Y, Tang D, Du Y, Liu B. Controlled Fabrication of Hexagonally Close-Packed Langmuir–Blodgett Silica Particulate Monolayers from Binary Surfactant and Solvent Systems. Langmuir [Internet]. 2013 Mar 5;29(9):2849–58. Available from: https://pubs.acs.org/doi/10.1021/la3049218

18. Vorobiev A, Paracini N, Cárdenas M, Wolff M. Π-GISANS: probing lateral structures with a fan shaped beam. Sci Rep [Internet]. 2021;11(1):1–8. Available from: https://doi.org/10.1038/s41598-021-97112-x

19. Müller-Buschbaum P. Grazing incidence small-angle neutron scattering: challenges and possibilities. Polym J [Internet]. 2013;45:34–42. Available from: https://www.nature.com/pj

20. Siffalovic P, Majkova E, Jergel M, Vegso K, Weis M, Luby S. Self-Assembly of Nanoparticles at Solid and Liquid Surfaces. In: Smart Nanoparticles Technology [Internet]. InTech; 2012. Available from: http://www.intechopen.com/books/smart-nanoparticles-technology/self-assembly-of-nanoparticles-at-solid-and-liquid-surfaces

21. Hrubý J, Santana VT, Kostiuk D, Bouček M, Lenz S, Kern M, et al. A graphene-based hybrid material with quantum bits prepared by the double Langmuir–Schaefer method. RSC Adv [Internet]. 2019;9(42):24066–73. Available from: http://xlink.rsc.org/?DOI=C9RA04537F

22. Pospelov G, Van Herck W, Burle J, Carmona Loaiza JM, Durniak C, Fisher JM, et al. BornAgain : software for simulating and fitting grazing-incidence small-angle scattering. J Appl Crystallogr [Internet]. 2020 Feb 1;53(1):262–76. Available from: http://scripts.iucr.org/cgi-bin/paper?S1600576719016789

23. Steinmetz NF, Bock E, Richter RP, Spatz JP, Lomonossoff GP, Evans DJ. Assembly of Multilayer Arrays of Viral Nanoparticles via Biospecific Recognition: A Quartz Crystal Microbalance with Dissipation Monitoring Study. Biomacromolecules [Internet]. 2008 Feb 1;9(2):456–62. Available from: https://pubs.acs.org/doi/10.1021/bm700797b

24. Luchini A, Nzulumike ANO, Lind TK, Nylander T, Barker R, Arleth L, et al. Towards biomimics of cell membranes: Structural effect of phosphatidylinositol triphosphate (PIP3) on a lipid bilayer. Colloids Surfaces B Biointerfaces [Internet]. 2019 Jan;17(August 2018):202–9. Available from: https://linkinghub.elsevier.com/retrieve/pii/S0927776518306428

25. Åkesson A, Lind T, Ehrlich N, Stamou D, Wacklin H, Cárdenas M. Composition and structure of mixed phospholipid supported bilayers formed by POPC and DPPC. Soft Matter [Internet]. 2012;8(20):5658. Available from: http://xlink.rsc.org/?DOI=c2sm00013j

26. Perrichon A, Devishvili A, Komander K, Pálsson GK, Vorobiev A, Lavén R, et al. Resonant enhancement of grazing incidence neutron scattering for the characterization of thin films. Phys Rev B [Internet]. 2021 Jun 17;103(23):235423. Available from: https://link.aps.org/doi/10.1103/PhysRevB.103.235423

27. Hafner A, Gutfreund P, Toperverg BP, Jones AOF, de Silva JP, Wildes A, et al. Combined specular and off-specular reflectometry: elucidating the complex structure of soft buried interfaces. J Appl Crystallogr [Internet]. 2021 Jun 1;54(3):924–48. Available from: https://doi.org/10.1107/S1600576721003575

28. Savarala S, Ahmed S, Ilies MA, Wunder SL. Formation and Colloidal Stability of DMPC Supported Lipid Bilayers on SiO 2 Nanobeads. Langmuir [Internet]. 2010 Jul 20;26(14):12081–8. Available from: https://pubs.acs.org/sharingguidelines

29. Drazenovic J, Ahmed S, Tuzinkiewicz N-M, Wunder SL. Lipid Exchange and Transfer on Nanoparticle Supported Lipid Bilayers: Effect of Defects, Ionic Strength, and Size. Langmuir [Internet]. 2015 Jan 20;31(2):721–31. Available from: https://pubs.acs.org/sharingguidelines

30. Ahmed S, Nikolov Z, Wunder SL. Effect of Curvature on Nanoparticle Supported Lipid Bilayers Investigated by Raman Spectroscopy. J Phys Chem B [Internet]. 2011 Nov 17;115(45):13181–90. Available from: https://pubs.acs.org/sharingguidelines

31. Tanaka M, Komikawa T, Yanai K, Okochi M. Proteomic Exploration of Membrane Curvature Sensors Using a Series of Spherical Supported Lipid Bilayers. Anal Chem [Internet]. 2020 Dec 15;92(24):16197–203. Available from: https://pubs.acs.org/doi/10.1021/acs.analchem.0c04039

32. Luchini A, Vitiello G. Understanding the Nano-bio Interfaces: Lipid-Coatings for Inorganic Nanoparticles as Promising Strategy for Biomedical Applications. Front Chem [Internet]. 2019 May 15;7:343. Available from: https://www.frontiersin.org

33. Brozell AM, Muha MA, Sanii B, Parikh AN. A Class of Supported Membranes: Formation of Fluid Phospholipid Bilayers on Photonic Band Gap Colloidal Crystals. J Am Chem Soc [Internet]. 2006 Jan 1;128(1):62–3. Available from: https://pubs.acs.org/doi/10.1021/ja056701j

