## Supplementary material for "Structural Characterisation of Nanoparticle-Supported Lipid Bilayers by Grazing Incidence X-ray and Neutron Scattering"

**Supplementary information and method section for:**
**Structural Characterization of Nanoparticle-Supported Lipid Bilayers by Grazing Incidence X-ray and Neutron Scattering**

### Materials:

1-palmitoyl-2-oleoyl-sn-glycero-3-phosphocholine (POPC) received from Larodan (Sweden), tail deuterated ($d_{64}$) POPC was produced by the National Deuteration Facility of the Australian Nuclear Science and Technology Organisation (ANSTO, Australia) and purified as previously described (1). Non-porous silica nanoparticles (NP) of 50 nm, 100 nm, 200 nm and 400 nm nominal diameter suspended in milliQ water were sourced from Alpha Nanotech (Canada). Deuterium oxide ($D_2O$) and cetyltrimethylammonium bromide (CTAB) were from Sigma Aldrich (Merk, Germany). The fluorescent lipid dye 1,2-dioleoyl-sn-glycero-3-phosphoethanolamine-N-(lissamine rhodamine B sulfonyl) (18:1 Liss Rhod PE) was from Sigma Aldrich (Merk, Germany). Silicon crystals (111) 60x80x10 mm, polished to a rms roughness <5 Å were from Sil'tronix (France). Glass slides for fluorescence microscopy with a diameter of 25 mm and a thickness of 0.16-0.19 mm were from Paul Marienfeld GmbH & Co (Germany) and were enclosed in Attofluor cell chamber from Invitrogen (Thermo Fisher Scientific inc. USA). All chemicals were used without further purification.

### Methods:

**Langmuir-Schaefer assembly of NP arrays**:

**NP preparation:** Silica NP were first exchanged from milliQ water into ethanol by centrifugation (10.000 rpm for 15 min) and resuspension (vortexing and sonication) until suspended in >99% ethanol according to manufacturer indications. After extensive vortexing and sonication in a cold bath, a 10 mg/ml suspension of particles was mixed 1:1 with a 2 mM ethanolic solution of CTAB to obtain a final concentration of 5 mg/ml NP and 1 mM CTAB in ethanol.

**Trough set-up:** A substrate (silicon crystal for scattering experiments, QCMD sensor, or glass slide fluorescent microscopy experiments) was cleaned by sonication in 2% SDS (10 min) followed by rinsing under milliQ water and sonication in hot ethanol (20 min), dried under $N_2$ and placed in a UV/ozone cleaner for 20 minutes. The substrate was then rinsed thoroughly under milliQ water, dried with $N_2$ and placed face-up on a holder inside a thoroughly cleaned custom-built Teflon Langmuir trough placed inside a fumehood (total trough surface 380x100 mm). The trough was filled with milliQ water until the water completely submerged the substrate's surface. The water surface was cleaned by aspiration with a nozzle connected to a pump. The water level was then adjusted by adding or removing water from behind the Teflon barrier so that the water level was only a few millimetres above the silicon crystal.

**NP isotherms and deposition**: Prior to spreading the NP onto the water surface, the 5 mg/ml NP, 1 mM CTAB ethanolic suspension was sonicated in a cold bath for 30 min. For the different NP sizes, volumes to deposit were calculate by estimating the number of NP required to yield a monolayer with an area corresponding to ~50% of the trough surface. The suspension was then deposited in a drop-wise manner onto the water surface using a 200 uL pipette from a height < 1 cm. The monolayer was left equilibrating for 15 min and then compressed to the target surface pressure (between 5-10 mN/m for all monolayers). Once the surface pressure stabilised, the compression was stopped, and the water level was lowered by slowly removing the subphase with a serological pipette connected to a pump from behind the barrier to avoid perturbing the monolayer until the substrate was completely emerged. The substrates were then left untouched inside the fume hood until dry.

**AFM:** A commercial Atomic Force Microscopy setup (MultiMode 8 SPM with a NanoScope V control unit, Bruker AXS, Santa Barbara CA) was used for imaging the transferred NP monolayer on the silicon wafer. Images were acquired by operating the AFM using PeakForce Tapping mode. For imaging in the PeakForce Tapping mode, cantilevers with a nominal resonance frequency between 320 and 364 kHz were used (RTESP7, Veeco Probes, Camarillo, CA).

**Fluorescence Microscopy:**
For super-resolution microscopy, images were acquired on a spinning disk system (Gataca Systems) based on a Nikon Ti2-E inverted microscope which also equipped with a super resolution module (Live-SR; Gataca systems) based on structured illumination with optical reassignment technique and online processing. The microscope was equipped with a sCMOS camera (Orca-Fusion; Hamamatsu), a confocal spinning head (W1; Yokogawa), a 100x 1.45 NA Plan-Apo objective lens. TIRF microscopy and FRAP images were acquired on the same microscope using iLAS ring-TIRF or FRAP system (GATACA Systems). All images were acquired using MetaMorph software (Molecular Devices, LLC, Sunnyvale, CA).

**GISAXS measurements:** GISAXS measurements were performed on a XEUSS 3.0 (Xenocs, France) equipped with a CuKα source (wavelength 1.54 Å) and a Pilatus 300K detector (Dectris, Switzerland) placed at 1700 mm from the sample. The beam was collimated in ultra-high resolution mode using slits of 300 x 150 μm and the angle of incidence was 0.2°. Samples assembled on silicon crystals, were measured for 3 hours in ambient conditions.

**GISAXS simulations:** GISAXS simulations were performed using the BornAgain software (2). A virtual instrument was created replicating the specifics used for data collection and described above as well as a background of Poisson noise. A model was set up reproducing a layer of silicon oxide spheres in a hexagonal lattice on a 10 Å thick silicon oxide layer (X-ray SLD 22.7 e-6 $Å^{-2}$) supported by an infinitely thick silicon substrate (X-ray SLD 20.0 e-6 $Å^{-2}$). The computational method employed to generate the images was the Monte Carlo integration. Parameters used in the simulations are given in **Figure S2**

**Neutron Reflectometry and GISANS**:

**Sample preparation**: Silicon crystals coated with the NP arrays were placed in the UV-ozone cleaner for 5 min, rinsed with milliQ water and dried under $N_2$, prior to assembly into solid liquid cells. POPC vesicles were prepared by thin film hydration and sonication in milliQ water at a concentration of 0.2 mg/ml. Immediately prior to vesicles injection in the cells the POPC suspension was diluted 1:1 with a 4 mM $CaCl_2$ solution to yield a final concentration of POPC 0.1 mg/ml and 2 mM $CaCl_2$. Vesicles were injected using a syringe pump at a flow rate of 1 ml/min. The injected vesicles were collected from the solid-liquid cell outlet and injected again in the opposite direction to maximise the lipid deposition.

**NR Measurements**: Neutron reflectometry measurements were carried out on the Figaro beamline at the Institute Laue Langevin (Grenoble)(3). Measurements were performed using a white neutron beam with wavelengths between 2 and 20 Å at two angles of incidence for the NP arrays (1° and 3.2°). For the NP-SLBs only the 1° angle was measured due to time constrains. Specular and off-specular signals were recorded on the area detector. $H_2O$ and $D_2O$ were flushed through the cells at 1 ml/min for 900 s to exchange solutions and EtOH washes were performed for 1200 s at the same flow rate.

**Specular NR data analysis**: Neutron reflectometry curves were fitted using the Rascal software (https://sourceforge.net/projects/rscl/). The reflectometry datasets were fitted to a mathematical model of the interface describing a silicon substrate covered with a layer of silicon oxide on top of which a monolayer of spheres was built from 2000 stacked thin slices, each one characterised by a defined thickness, SLD and roughness, further described in **Figure S1** and **S4**. For the 50 nm and 100 nm NP arrays, two datasets per sample (collected in $H_2O$ and in $D_2O$) were constrained to fit to a common structure of the monolayer, only allowing the SLD of the solvent to vary. For the 200 nm NP used to assemble the NP-SLB, the model was expanded to fit together six datasets which all shared the same structure of the spheres (i.e. thickness and spheres volume fraction). Two datasets were the NP array in $H_2O$ and $D_2O$, two the NP-SLB assembled using hPOPC and two the NP-SLB assembled using tail deuterated $d_{64}$POPC. Data were fitted by using a simplex algorithm to minimise the difference between model and data (chi$^2$). Error analysis was performed using the bootstrap error analysis built into the software (100 runs 3000 iterations per run) which resamples the data points and repeats the fits starting from random parameters.

**Off-specular NR analysis**: Quantitative fits of the off-specular scattering patterns were performed using the algorithm described in (4). The specular reflectivity model consisting of 2000 smooth equidistant slabs was coarse-grained to 7 layers in case of bare NPs and 9 layers in case of NP-SLBs of variable thickness and roughness reproducing closely the high-resolution SLD profile of the specular model. The top 6 slabs from this coarse-grained SLD profile, which correspond to the NP layer, are assumed to include cylindrical in-plane inhomogeneities with SLDs corresponding to $SiO_2$ ($3.47 \times 10^{-6}$Å$^{-2}$) and $D_2O$ ($6.35 \times 10^{-6}$Å$^{-2}$), respectively, of variable radius corresponding to a sphere form factor. The positions of these cylinders were assumed to be out-of-plane correlated within these 6 slabs. The volume fraction of the respective cylinders are taken such that the mean SLD of the layer matches the SLD from the specular fits. Once the (protonated) lipid is added, two models gave equally good fits to the data starting from the fixed parameters from the bare NPs in $D_2O$: in the first case, the lipids are spread homogeneously between the NPs (as if they were in solution) and in the second a third phase (inhomogeneity) is present in between the $SiO_2$ cylinders and has a thickness and SLD of a hydrogenous lipid bilayer. Although both models yielded the same off-specular scattering patterns, only the latter model is physically relevant and in agreement with the results from the other techniques.

**GISANS measurements**: GISANS measurements were performed on D22 at the Institute Laue Langevin using a monochromatic beam with a wavelength of 6 Å. The sample was illuminated for 3 h at an angle of 0.35° and the area detector was placed 17600 mm away from the sample stage.

**QCMD measurements**: QCMD measurements were performed on a QSense Analyzer (Biolin Scientific, Sweden). Of the 4 flow modules two were equipped with flat silicon oxide sensors and two with sensors coated with a 200 nm NP array and were left equilibrating in water until the frequency and dissipation signals stabilised. Solutions were flushed through the cells using a peristaltic pump at a speed of 0.1 ml/min. Frequency and dissipation shifts of the 7th harmonic were measured for the lipid deposition and removal processes. Values reported in the text are the average and standard deviation of 4 lipid depositions on two different sensors (two deposition on each sensor).

### Supplementary figures:

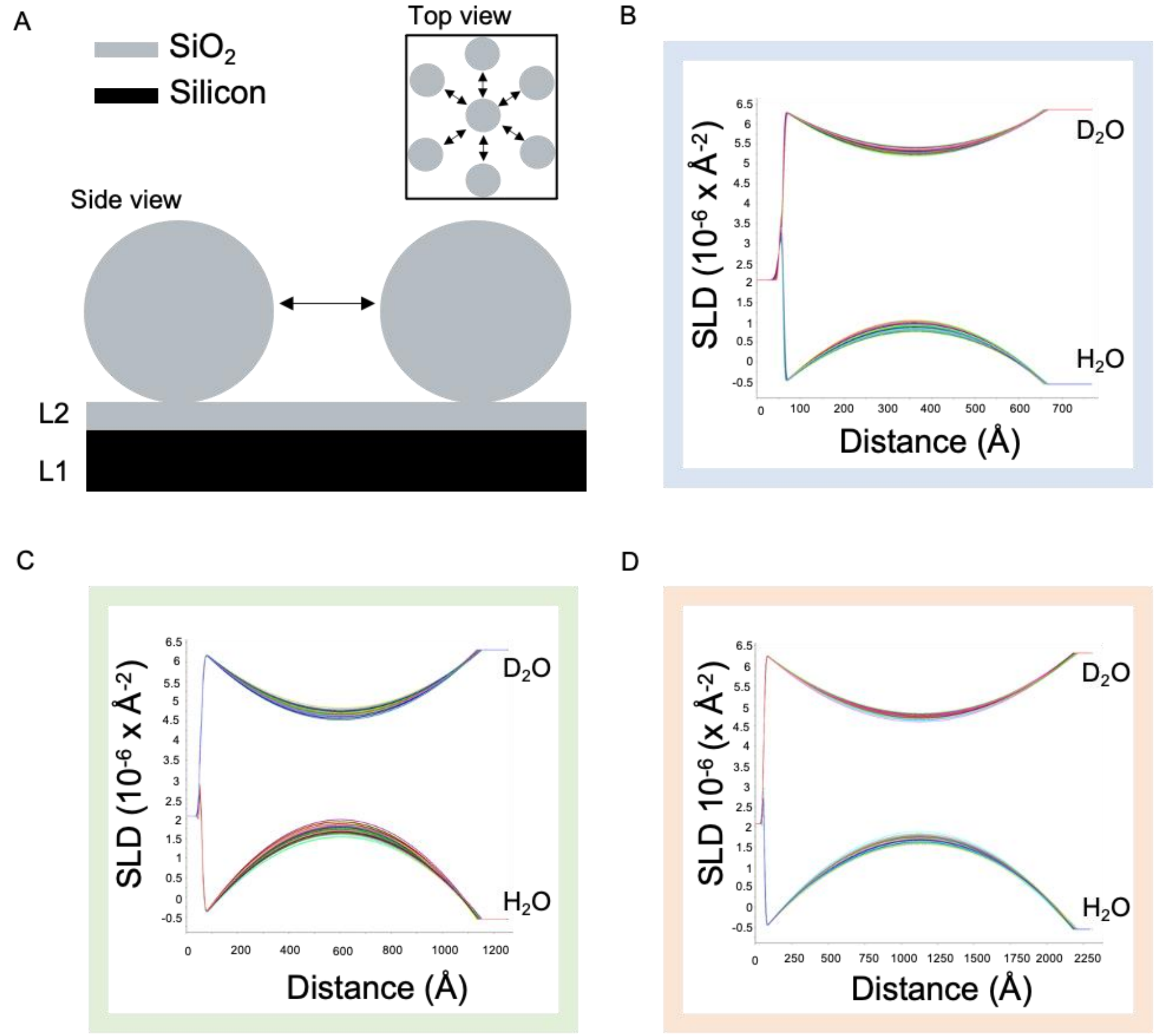


**Figure S1 – Analysis of the reflectivity data from the NP monolayers in water.** (**A**) Schematic representation of the model used to fit the reflectivity data from the NP monolayers composed of the silicon substrate (black, labelled L1), a thin layer of $SiO_2$ (grey, labelled L2), and the $SiO_2$ NP (grey). The coverage parameter determines the average distance between the spheres indicated by the black double pointed arrows. (**B, C** and **D**) Bootstrap error analysis obtained from resampling and fitting the NR data collected in $D_2O$ and $H_2O$ (shown in **Figure 2A**) to a common model of the interface for the three NP monolayers of increasing size. Nominal diameters are 500 Å, 1000 Å and 2000 Å in **B**, **C** and **D** respectively

**Table S1 – Parameters relative to the fits of the NR data from the three NP monolayers**
Parameters obtained from the fits shown in **Figure 2A** and **B** and the error analysis shown in **Figure S1,** color coding refers to the respective fits in Fig S1.
[a]Parameters fixed to the calculated value

| | Small size NP | | Medium size NP | | Large size NP | |
|---|---|---|---|---|---|---|
| **Parameter** | **Estimated value** | **Estimated error** | **Estimated value** | **Estimated error** | **Estimated value** | **Estimated error** |
| L1 roughness (Å) | 2.5 | ± 2.3 | 2.0 | ± 1.6 | 7.5 | ± 0.6 |
| L2 thickness (Å) | 11.5 | ± 0.7 | 11.2 | ± 0.5 | 10.6 | ± 0.2 |
| L2 roughness (Å) | 3.0 | ± 0.6 | 7.5 | ± 0.3 | 6.2 | ± 2.8 |
| L2 hydration (%) | 0.3 | ± 0.1 | 0.3 | ± 0.1 | 1.2 | ± 0.4 |
| Sphere diameter (Å) | 604.0 | ± 4.6 | 1085.6 | ± 8.3 | 2148.8 | ± 12.1 |
| Sphere coverage (%) | 40.4 | ± 4.2 | 62.0 | ± 5.0 | 66.2 | ± 1.4 |
| $SiO_2$ SLD | 3.47e-6[a] | - | 3.47e-6[a] | - | 3.47e-6[a] | - |
| Silicon SLD | 2.07e-6[a] | - | 2.07e-6[a] | - | 2.07e-6[a] | - |

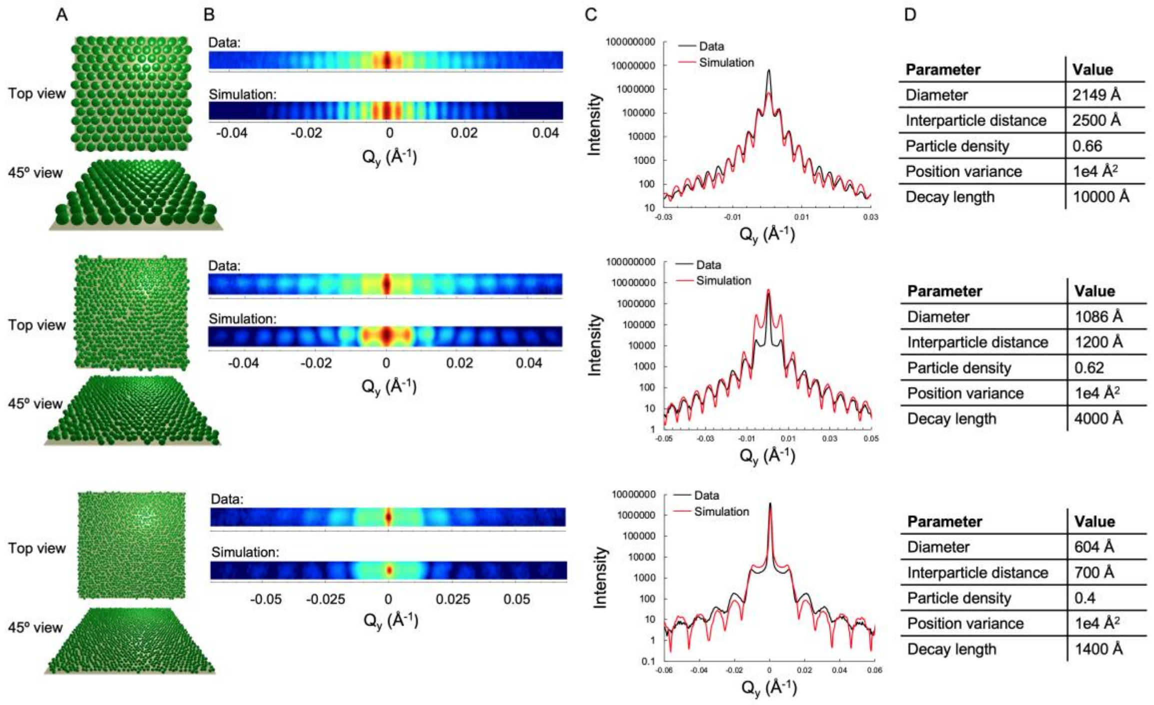


**Figure S2 Comparison between GISAXS data and simulations** (**A**) 3D views of the particle layouts that generated the GISAXS signals in the simulation runs with BornAgain. (**B**) Data (top) and simulations (bottom) of GISAXS signal around the specular peak (**C**) Comparison of the integrated signals shown in **B**, simulations shown in red and data in back (**D**) Values of the parameters used to generate the simulated GISAXS profiles. The decay length represents the full-width half maximum of the Cauchy 2D distribution function.

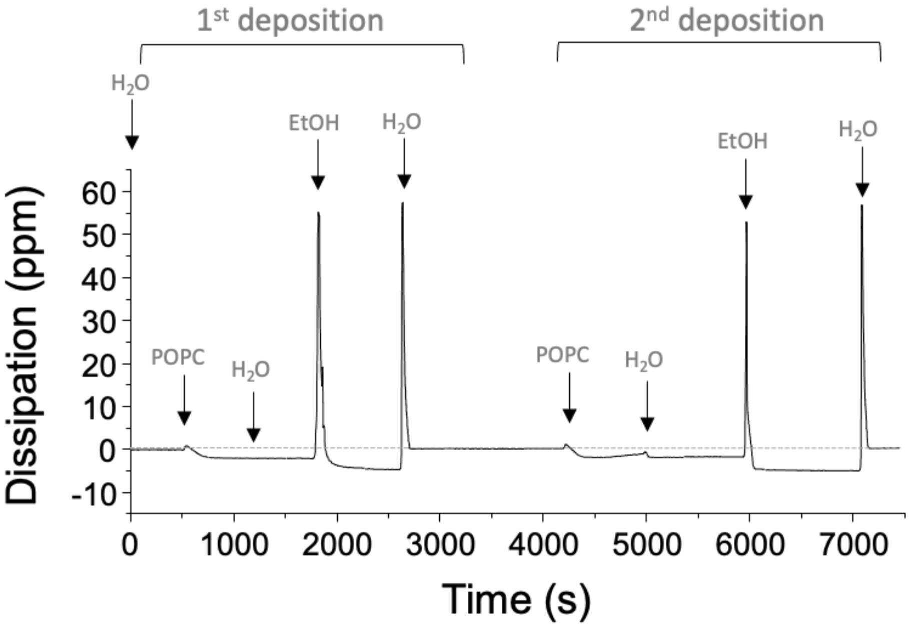


**Figure S3 QCMD dissipation trace relative to the frequency shifts shown in figure 3A**

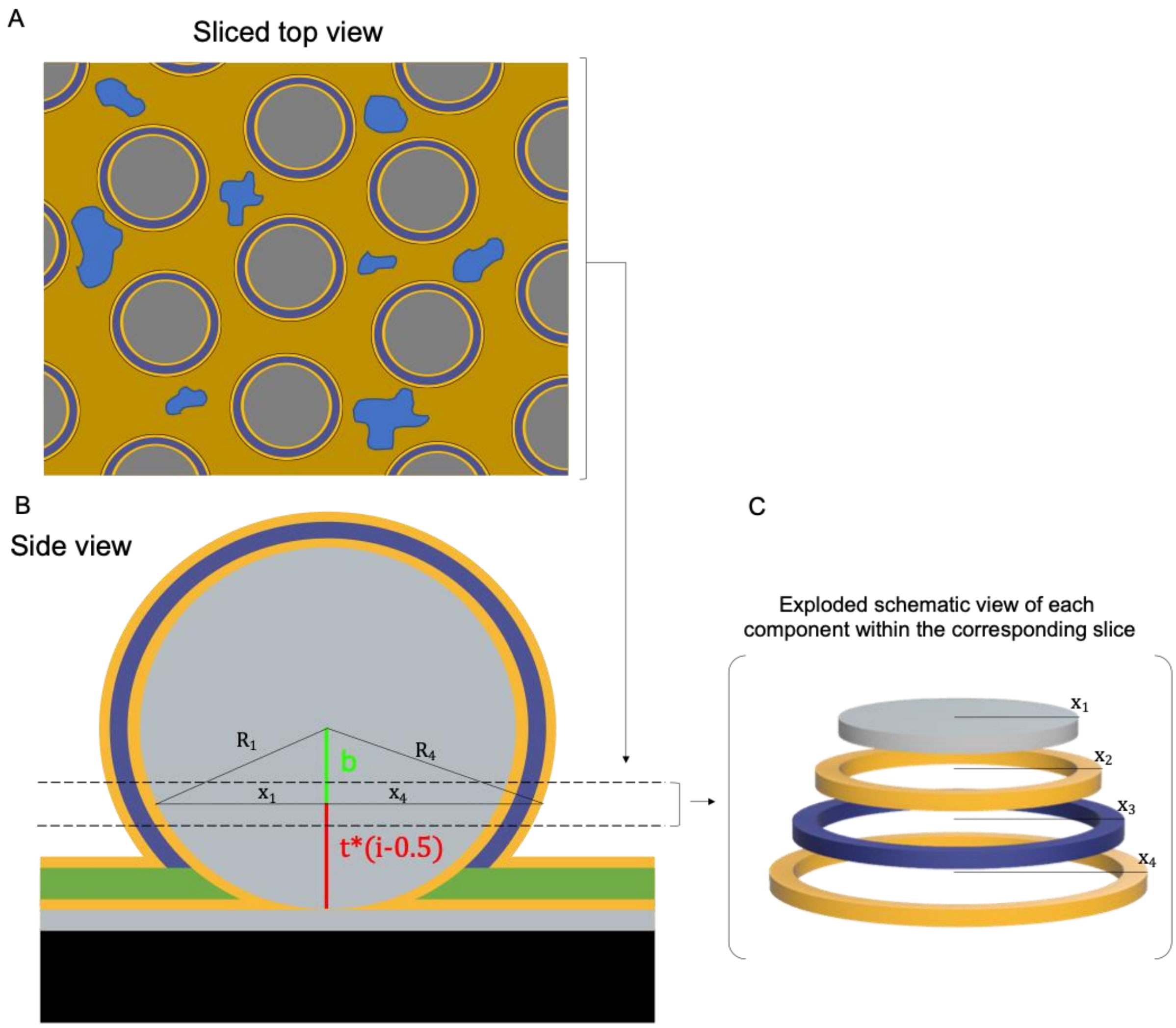


**Figure S4 Modelling of the NR data from the NP-SLB.** (**A**) Illustration of an array of lipid-coated nanoparticles (top view, sliced along the dashed line in **B**). The blue areas represent incomplete coverage of the flat lipid bilayer formed on the surface below the NP. (**B**) Side view of a lipid coated NP (thickness of the different layers not to scale). (**C**) Composition of a sample slice in the NP-SLP model.

To generate a model of the interface formed by the NP and the lipid coating as shown in the figure, we considered the volume contribution of the various components in slices away from the surface. For example, between the dashed lines in the side view, the contribution of the NP sphere and the lipid coating give a thin cylinder corresponding to the NP (grey) and three hollow cylinders corresponding to the inner and outer head group regions (yellow) and the tail region (purple), with the radii of these ($x_1$, $x_2$, $x_3$ and $x_4$) dependent on the distance from the surface.

The model was parametrized using the following mathematical description.

Given:

$t$ = thickness of each slice

$i$ = slice index

$R_1$ = radius of the sphere

If $t * i < R_1$ then $b = R_1 - t(i - 0.5)$ (i.e. up to the midpoint of the NP)

If $t * i > R_1$ then $b = -(R_1 - t(i - 0.5))$ (i.e. above the midpoint of the NP)

Therefore $b^2 = \left(R_1 - t(i - 0.5)\right)^2$ across the whole diagram, and by Pythagoras:

$x_1^2 = R_1^2 - b^2$

$x_2^2 = R_2^2 - b^2$

$x_3^2 = R_3^2 - b^2$

$x_4^{\ 2} = R_4^{\ 2} - b^2$

Where:

$R_2 = R_1 +$ inner headgroup thickness

$R_3 = R_2 +$ tails thickness

$R_4 = R_3 +$ outer headgroup thickness

The volume fraction of the materials within each slice was calculated by adding the volumes of individual components and dividing the total by the volume of the slice, which was calculated assuming hexagonal packing of the spheres. The volume fraction of the slice that was not occupied by the NP nor the lipids, was occupied by the solvent. An additional coverage parameter was used to take into account defects of the hexagonal packing by allowing additional solvent to hydrate empty space between the coated speres. The model used to fit NP in the absence of lipids was identical but did not include any coating of the spheres.

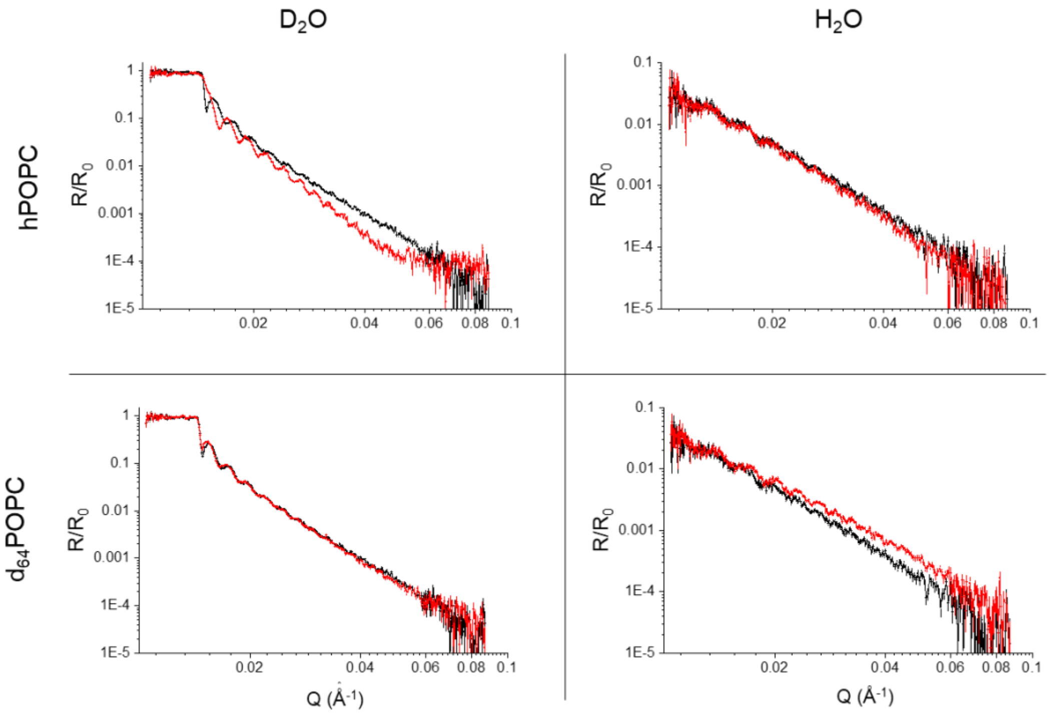


**Figure S5 Effect of lipid addition on reflectivity profiles at different isotopic contrasts** NR data before (black) and after (red) lipid addition to the NP array in $H_2O$ and $D_2O$.

**Table S2 – Parameters relative to the fits of the NR data describing the hydrogenous and deuterated NP-SLB**

The parameters describing the NP array and the underlying substrate are shown in the “large size NP” section of table **Table S1.** The terms hydration and coverage can be used interchangeably as they describe complementary quantities that add up to 1 (i.e. 35% hydration = 65% coverage of a certain component). Tails thickness values refer to the thickness of the entire hydrophobic core of the bilayer.

[a]Parameters fixed to the calculated value

[b]Headgroups hydration values refer only to the water associated with the headgroups. The total volume fraction in the head group layers amounts to the sum of the head and tails hydration.

| | Parameter | Value | Estimated error |
|---|---|---|---|
| **Shared by both bilayers** | POPC heads SLD ($Å^{-2}$) | 1.8e-6[a] | - |
| | hPOPC tails SLD ($Å^{-2}$) | -3.9e-7[a] | - |
| | dPOPC tails SLD ($Å^{-2}$) | 6.1e-6 | ± 1.2e-7 |
| **Bilayer on the flat surface** | POPC heads thickness (Å) | 5.6 | ± 1.3 |
| | POPC tails thickness (Å) | 32.2 | ± 2.1 |
| | hPOPC heads hydration (%) | 42.8[b] | ± 1.6 |
| | hPOPC tails hydration (%) | 39.4 | ± 3.2 |
| | dPOPC heads hydration (%) | 44.3[b] | ± 1.1 |
| | dPOPC tails hydration (%) | 38.6 | ± 0.5 |
| **Bilayer coating the NP** | POPC heads thickenss (Å) | 6.0 | ± 1.1 |
| | POPC tails thickness (Å) | 27.5 | ± 1.5 |
| | hPOPC heads hydration (%) | 60.00[b] | ± 1.4 |
| | hPOPC tails hydration (%) | 1.2 | ± 0.7 |
| | dPOPC heads hydration (%) | 61.8[b] | ± 2.0 |
| | dPOPC tails hydration (%) | 3.3 | ± 1.9 |

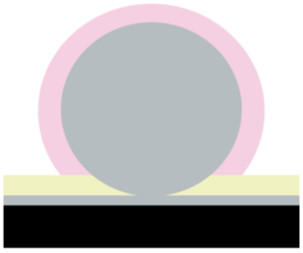

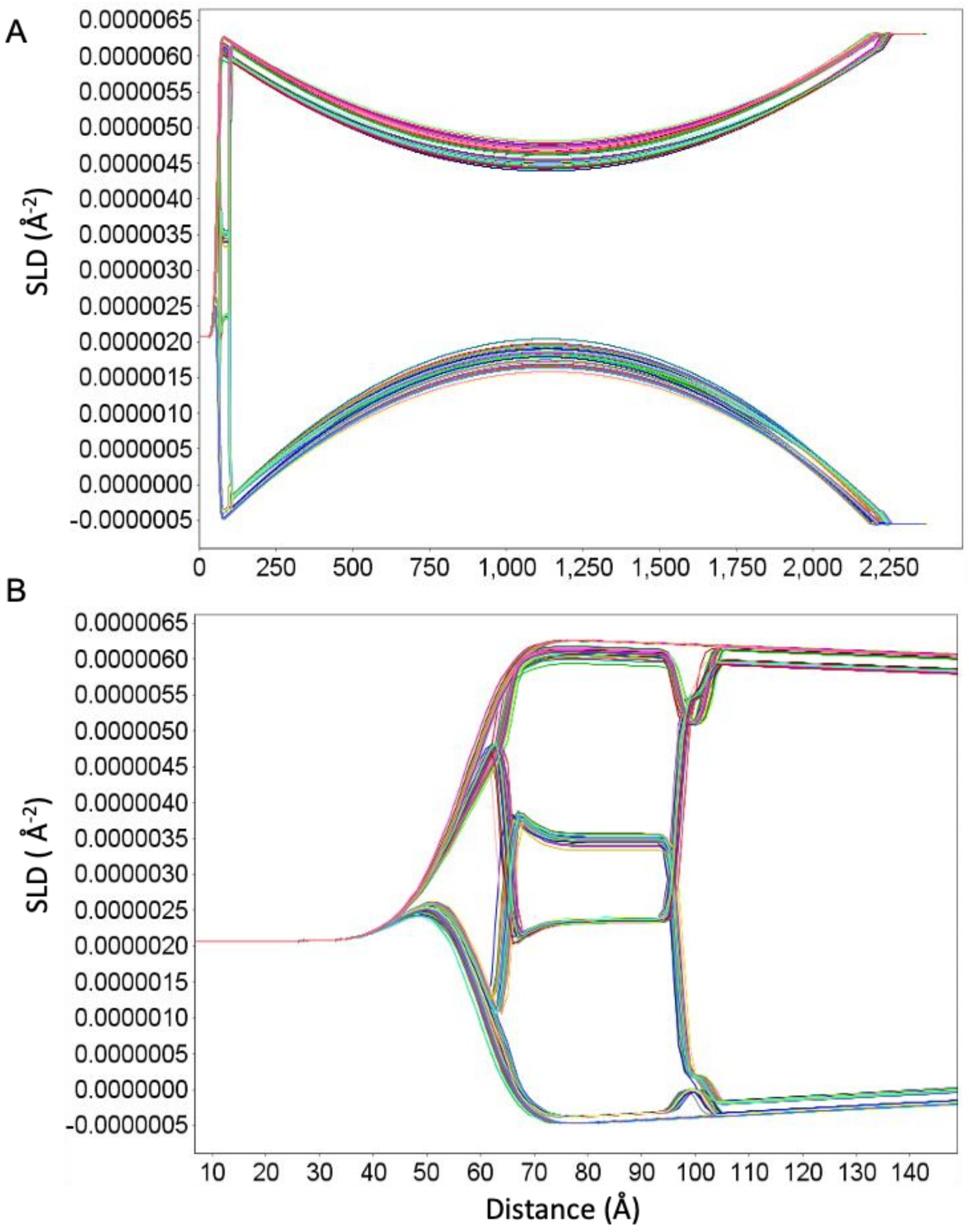


**Figure S6 – Bootstrap error analysis of the NR data collected on the NP-SLB.** (**A**) Analysis of the SLD profiles obtained from resampling and fitting six NR curves to a common model of the interface. The six data sets are the NP monolayer in $H_2O$ and in $D_2O$, hPOPC NP-SLB in $H_2O$ and in $D_2O$ and $d_{64}$POPC NP-SLB in $H_2O$ and in $D_2O$. (**B**) close up of the region close to the interface showing the planar SLB formed on the silicon crystal surface.

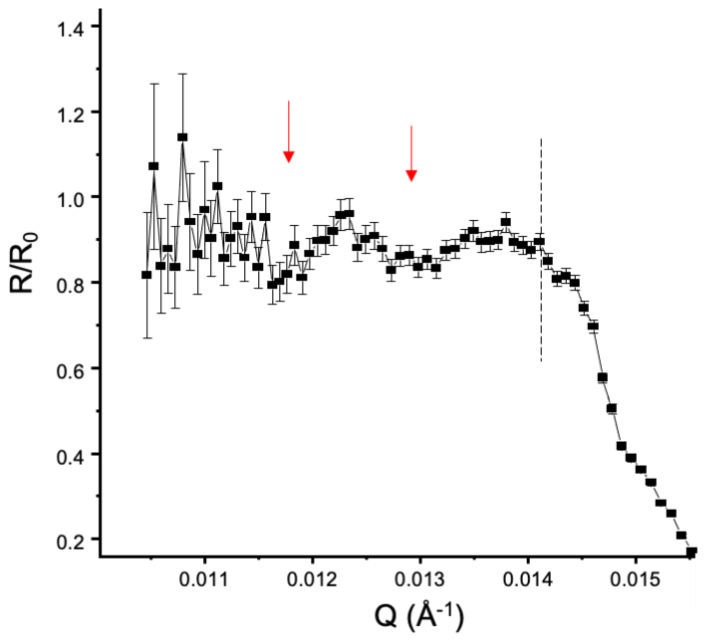


**Figure S7 Intensity dips due to the resonances below the critical Q** Reflectivity measured on the hPOPC NP-SLB in $D_2O$**.** The dashed line indicates the critical edge below which total reflection occurs.

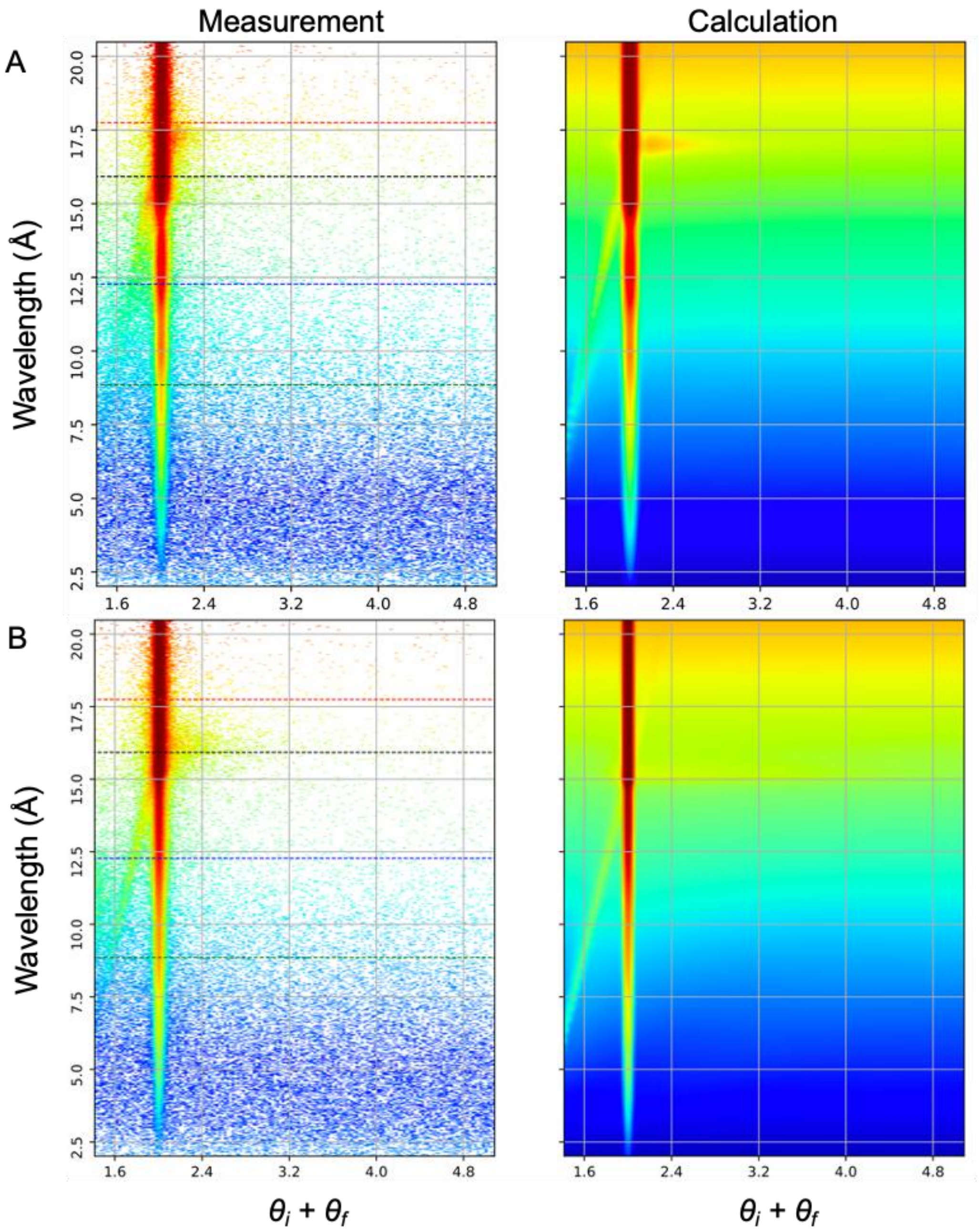


**Figure S8 Measured and simulated off-specular signal** Off-specular reflectometry signals (left) and simulated image (right) for 100 nm (**A**) and 50 nm (**B**) NP monolayers in $D_2O$.

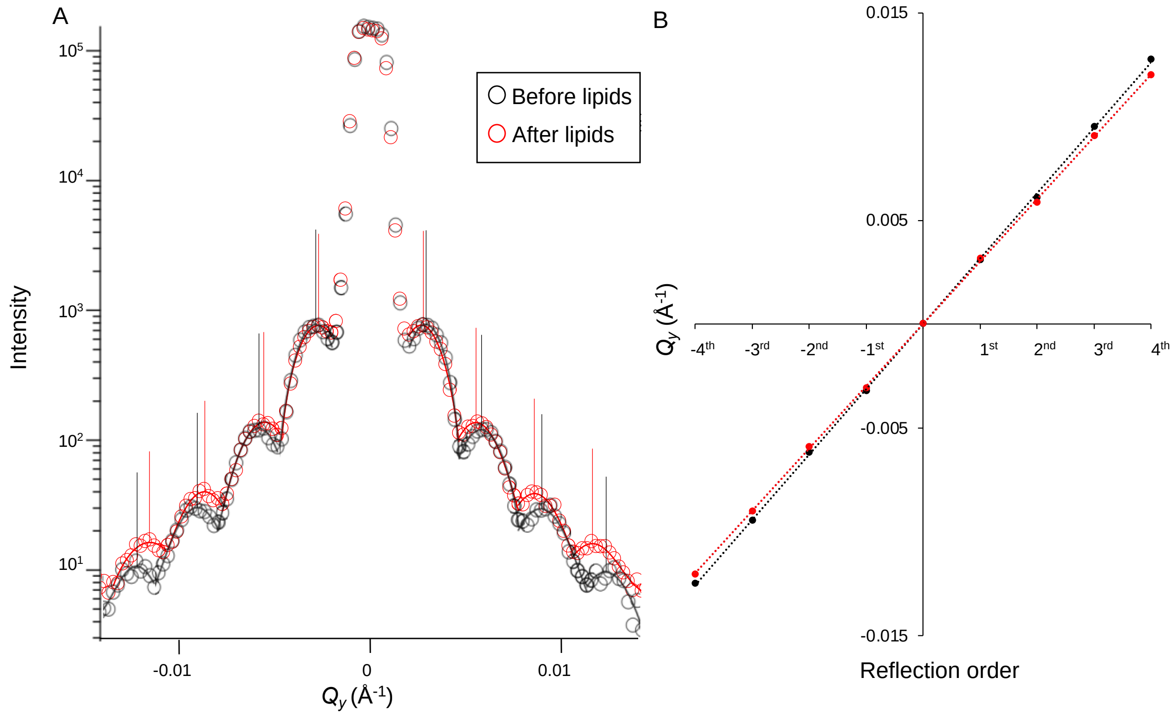


**Figure S9 Analysis of the GISANS peak positions** (**A**) Gaussian fits and labelled maxima of individual GISANS peaks of the 200 nm NP array before (black) and after (red) addition of hPOPC (**B**) Linear regression of the peak positions.


1. Waldie S, Sebastiani F, Browning K, Maric S, Lind TK, Yepuri N, et al. Lipoprotein ability to exchange and remove lipids from model membranes as a function of fatty acid saturation and presence of cholesterol. Biochim Biophys Acta - Mol Cell Biol Lipids [Internet]. 2020 Oct;1865(10):158769. Available from: https://doi.org/10.1016/j.bbalip.2020.158769
2. Pospelov G, Van Herck W, Burle J, Carmona Loaiza JM, Durniak C, Fisher JM, et al. BornAgain : software for simulating and fitting grazing-incidence small-angle scattering. J Appl Crystallogr [Internet]. 2020 Feb 1;53(1):262–76. Available from: http://scripts.iucr.org/cgi-bin/paper?S1600576719016789
3. Campbell RA, Wacklin HP, Sutton I, Cubitt R, Fragneto G. FIGARO: The new horizontal neutron reflectometer at the ILL. Eur Phys J Plus [Internet]. 2011 Nov 11;126(11):107. Available from: http://link.springer.com/10.1140/epjp/i2011-11107-8
4. Hafner A, Gutfreund P, Toperverg BP, Jones AOF, de Silva JP, Wildes A, et al. Combined specular and off-specular reflectometry: elucidating the complex structure of soft buried interfaces. J Appl Crystallogr [Internet]. 2021 Jun 1;54(3):924–48. Available from: https://doi.org/10.1107/S1600576721003575
